# Cholinergic Modulation is Necessary for Upward Firing Rate Homeostasis in Rodent Visual Cortex

**DOI:** 10.1101/2023.04.11.536412

**Authors:** Juliet Bottorff, Sydney Padgett, Gina G. Turrigiano

**Affiliations:** Brandeis University, Department of Biology, 415 South Street Waltham MA 02453, USA

## Abstract

Bidirectional homeostatic plasticity allows neurons and circuits to maintain stable firing in the face of developmental or learning-induced perturbations. In primary visual cortex (V1), upward firing rate homeostasis (FRH) only occurs during active wake (AW) and downward during sleep, but how this behavioral state-dependent gating is accomplished is unknown. Here we focus on how AW enables upward FRH in V1 of juvenile Long Evans rats. A major difference between quiet wake (QW) when upward FRH is absent, and AW when it is present, is increased cholinergic (ACh) tone; we therefore chemogenetically inhibited V1-projecting basal forebrain cholinergic (BF ACh) neurons while inducing upward FRH using visual deprivation, and found that upward FRH was completely abolished. Next, we examined the impact on synaptic scaling and intrinsic excitability, two important cellular targets of homeostatic regulation. BF ACh inhibition impaired synaptic scaling up, and dramatically decreased the intrinsic excitability of activity-deprived V1 pyramidal neurons, consistent with the block of upward FRH. Interestingly, knock down of the highly abundant M1 ACh receptor in V1 failed to phenocopy the effects of decreased BF ACh activity on intrinsic excitability, suggesting either that BF ACh activity acts through a different receptor within V1, or acts indirectly via other brain regions or cell types. Together, our results show that BF ACh modulation is a key enabler of upward homeostatic plasticity, and more broadly suggest that neuromodulatory tone is a critical factor that segregates upward and downward homeostatic plasticity into distinct behavioral states.

**SIGNIFICANCE STATEMENT:** Hebbian, positive feedback-based and homeostatic, negative feedback-based plasticity mechanisms are necessary to maintain the functionality of flexible yet stable complex neuronal circuits. Growing evidence suggests a role for behavioral state in temporally segregating these opposing plasticity mechanisms, but how behavioral states enact this gating remains unknown. Here, we tested the role of acetylcholine (ACh), a widespread neuromodulator largely released during active wake, in the regulation of upward homeostatic plasticity. We found that ACh modulation is indeed necessary for the expression of active wake-gated upward firing rate homeostasis, likely due to its role in maintaining intrinsic excitability of cortical pyramidal neurons. Our results suggest potential mechanisms by which neuromodulatory tone may enable behavioral state gating of homeostatic plasticity.

## INTRODUCTION

Homeostatic plasticity compensates for developmental or learning-induced perturbations by globally and bidirectionally adjusting synaptic and neuronal properties to maintain average firing within a ‘set point’ range (firing rate homeostasis: Turrigiano and Nelson, 2004; Turrigiano 2017; Hengen et al., 2013). In rodent monocular visual cortex (V1m), upward and downward firing rate homeostasis (FRH) occur during distinct behavioral states (Hengen et al., 2016; Torrado-Pacheco et al., 2021), but how this state-dependent segregation is achieved on a mechanistic level is unknown. Upward FRH occurs exclusively during active waking (AW) states when cholinergic inputs to cortex are most active, suggesting that these inputs might be essential for its induction.

During monocular deprivation (MD) in V1m, firing rates drop dramatically in the first two days of MD due to the induction of long-term depression (LTD)-like plasticity (Rittenhouse et al., 1999; Miska et al., 2018); this perturbation of firing then triggers homeostatic plasticity that slowly returns firing rates back to baseline during MD 3-4, even though the eye remains closed (Mrsic-Flogel et al., 2007; Kaneko et al., 2008; Hengen et al., 2013, 2016). This process is bidirectional, as reopening the eye after compensation has occurred induces a dramatic overshoot of firing that is dependent on an LTP-like process, which is again followed by a slow homeostatic return to baseline (Torrado Pacheco et al., 2021). Remarkably, both upward and downward FRH are gated by behavioral state, but in opposite directions: upward FRH only occurs during AW, while downward FRH only occurs during sleep (Hengen et al., 2016; Torrado Pacheco et al., 2021).

Sensory inputs and neural activity patterns are dramatically different during wake and sleep; however, the difference between quite wake (QW) when upward firing rate homeostasis is absent, and AW when it is robustly induced, is much subtler and largely due to differences in the activity of neuromodulatory systems (Steriade et al., 2001; Jones, 2005; Lee and Dan, 2012). In particular, increased cholinergic (ACh) modulation is a key signature of AW, and optogenetic stimulation of ACh neurons promotes rapid transitions from non-rapid eye movement sleep (NREM) to AW (Lee and Dan, 2012, Han et al., 2014; Zant et al., 2016). ACh inputs to rodent V1 come almost exclusively from the horizontal diagonal band (HDB) of the basal forebrain (BF); these neurons fire at high rates during AW and are almost silent during NREM and QW (Lee et al., 2005; Xu et al., 2015; Kim et al., 2016). Notably, ACh is known to be important for the expression of ocular dominance plasticity in V1 (Bear and Singer, 1986), which relies on homeostatic plasticity (Espinosa and Stryker, 2012), and to enhance multiple forms of visual learning (Pinto et al., 2014; Kang et al., 2015).

Here we test the hypothesis that activity of HDB ACh neurons is essential for enabling upward firing rate homeostasis. We use chemogenetics to chronically inhibit HDB ACh neurons during MD 3-4, without inducing major changes to the animals’ sleep/wake behavior. Strikingly, we find that this manipulation completely prevents the expression of upward firing rate homeostasis. To explore the cellular plasticity mechanisms that underlie this effect, we induced homeostatic compensation in V1 using local DREADDs-mediated inhibition of pyramidal neurons and found that reducing the activity of HDB ACh inputs occluded synaptic scaling. Further, HDB inputs were crucial for maintaining the intrinsic excitability of layer (L)2/3 pyramidal neurons during activity deprivation, suggesting that cholinergic regulation of intrinsic plasticity plays an important role in upward firing rate homeostasis. To test whether this regulation occurs locally within V1, we used a viral strategy to knock down expression of the abundant Gq-coupled M1 ACh receptor; while this paradigm dramatically reduced cholinergic modulation of L2/3 pyramidal neurons measured *ex vivo*, it did not phenocopy the effects of HDB inhibition, suggesting that the site of action may be external to V1. Taken together, our data show that ACh modulation plays a critical role in gating upward firing rate homeostasis, in large part through modulating the intrinsic excitability of V1 pyramidal neurons.

## MATERIALS AND METHODS

### Animals

Experiments were performed on Long Evans rats (both wild-type (WT) and transgenic rat lines) of both sexes. All ChAT-Cre transgenic rats (Witten et al., 2011) were bred as heterozygotes and continuously back-crossed with WT Long Evans rats. Experiments were carried out between postnatal day 24 (P24) and P32, during the classical visual system critical period. All animals were treated in accordance with Brandeis Institutional Biosafety Committee and Institutional Animal Care and Use Committee protocols. Pups were weaned at P21 and housed (12 hr L/D cycle, *ad libitum* access to food and water) with at least one littermate except for post-surgical recovery when experiments required single housing. The number of animals and neurons for each experiment are given in the figure legends, and individual data points represent neurons, except in Figure 1, where they represent animals.

**Figure 1:**
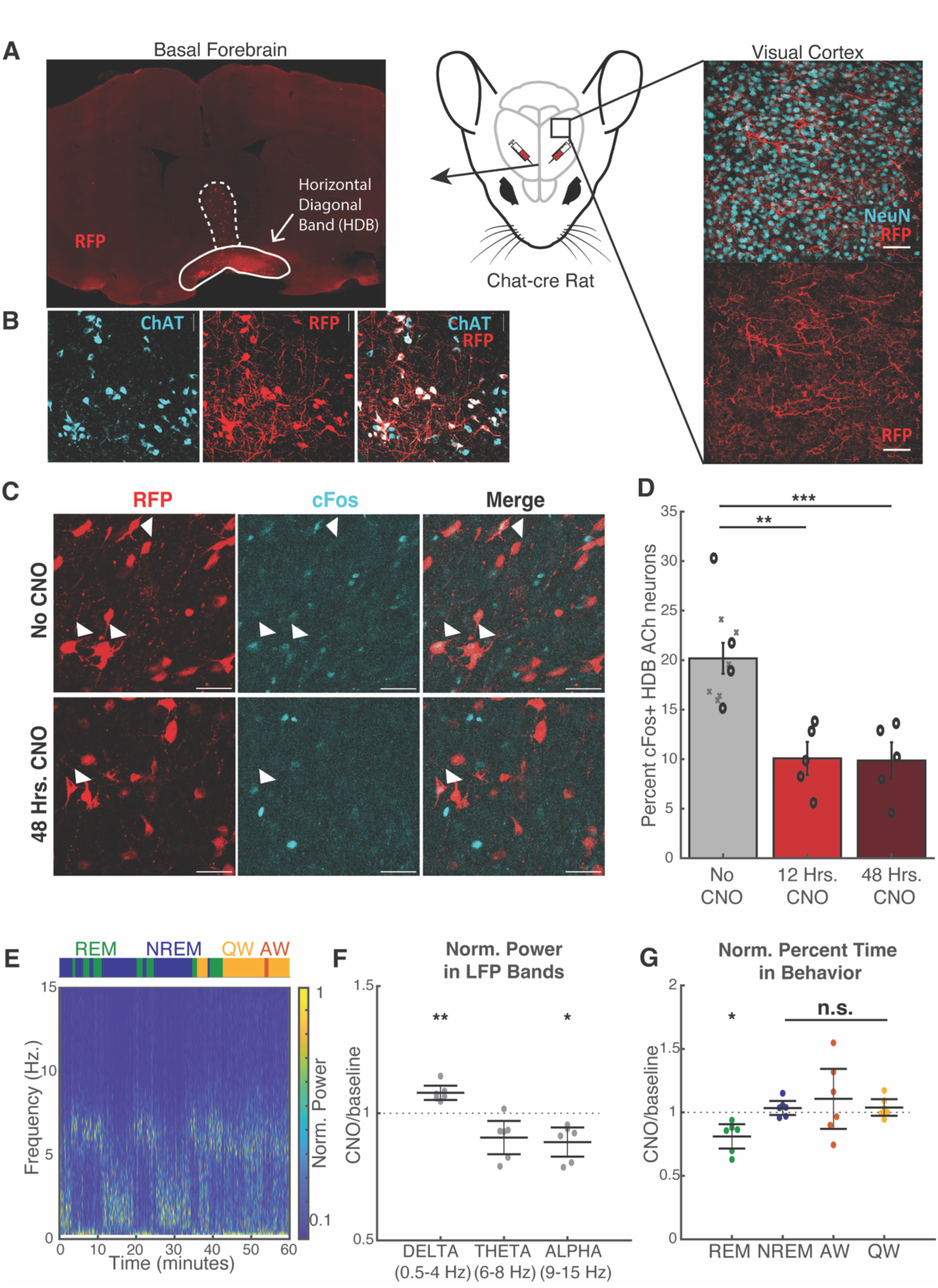
Long-lasting inhibition of V1-projecting HDB ACh neurons using DREADDs. A) Schematic of viral injection of AAV-hsyn-DIO-hM4D(Gi)-mCherry into HDB of ChAT-cre rats (Center; rat head adapted from Scidraw) and example images of the resulting mCherry+ cell bodies in the basal forebrain (Left) and their mCherry+ projections in visual cortex (Right). RFP labels the mCherry tag. **Left image** in basal forebrain: solid line indicates basal forebrain region of interest, the horizontal diagonal band (HDB); dashed line indicates vertical diagonal band, another region of the basal forebrain; 10x magnification; red = RFP. **Right image** in visual cortex: scale bar = 50 μm; cyan = NeuN; red = RFP. B) mCherry expression from virus injections is restricted to ChAT+ neurons in ChAT-cre rats. Scale bar = 50 μm; cyan = ChAT; red = RFP. C) Example images of cFos and mCherry expression after virus injections into the HDB of ChAT-cre rats. Top row is from an animal without any exposure to CNO, bottom row is from an animal that received CNO in its drinking water for 48 hours prior to sacrifice. Scale bar = 50 μm; cyan = cFos; red = RFP. D) Quantification of cFos expression in HDB ACh neurons, showing ∼50% reduction in cFos expression after either 12 or 48 hours of CNO exposure. All animals in the 12 hour CNO (N=5) and 48 hour CNO (N=5) groups had Hm4di-mCherry in BF ACh neurons, and quantification was evaluated as the percent of RFP+ HDB neurons that were cFos+. The No CNO group includes 4 animals with Hm4di-mCherry in HDB ACh neurons (black circles) and 6 animals without virus injection (gray X’s). The Hm4di-mCherry animals were quantified as above, and the no virus injection animals were quantified as the percent of ChAT+ HDB neurons that were cFos+. We note that there was no difference in these two No CNO groups, indicating no effect of Hm4di-mCherry on basal HDB ACh neuron activity, so we combined the two groups here. P = 0.00018, 1 way ANOVA with Tukey’s post hoc. No CNO vs. 12 Hrs. CNO: p = 0.0011; No CNO vs. 48 Hrs. CNO: p = 0.0009. E) Example LFP spectrogram and behavior from 1 hour of baseline recording in one rat. LFP was extracted from an electrode array in visual cortex, and color indicates power, normalized to the maximum value at each time point. Top: Colored boxes indicate the rat’s behavioral state over the same time course as the LFP spectrogram. Behavior was coded using LFP, EMG, and video data. Blue = Non-REM Sleep; green = REM Sleep; yellow = Quiet Wake; orange = Active Wake. F) Average power in each of three LFP frequency bands in the 8 hours after subcutaneous CNO injections, normalized to time- and circadian-matched baseline periods on recording days without CNO injections. Since each animal received 4 total CNO injections, spaced 12 hours apart, normalized values for each CNO injection were averaged per animal, so that each data point represents one animal. Dotted line at 1 represents no change before and after CNO injections (HDB ACh inhibition). 1 sample t-test comparing normalized values to no change, with Bonferroni correction for multiple comparisons. Delta = 0.5-4Hz; p = 0.0075. Theta = 6-8Hz; p = 0.11. Alpha = 9-15Hz; p = 0.035. G) Percent time spent in each of 4 behavioral states in the 8 hours after subcutaneous CNO injections, normalized to time- and circadian-matched baseline periods before the CNO injections. As above, each data point represents one animal. Dotted line at 1 represents no change. 1 sample ttest comparing each normalized value to no change, with Bonferroni correction for multiple comparisons. REM = Rapid Eye Movement Sleep; p = 0.047. NREM = Non-REM Sleep; p = 1. AW = Active Wake; p = 1. QW = Quiet Wake; p = 1. **F** and **G**: N = 6 ChAT-cre rats, all with Hm4di virus injections into the HDB.

### Virus Injection

Virus injections were performed on juvenile rats on a stereotaxic apparatus under isoflurane anesthesia. V1m or the horizontal diagonal band (HDB) region of the basal forebrain (2 locations per hemisphere) were targeted using the following stereotaxic coordinates determined from Paxinos and Watson (1998): for V1m, (1) A/P -7.3 - -6.9 mm; M/L ± 2 – 3 mm; D/V 0.8 – 0.4 mm; and (2) A/P -6.7 mm; M/L ± 2.8 – 3 mm; D/V 0.8 – 0.4 mm; for HDB, (1) A/P 0.3 mm; M/L ± 0.3 – 0.5 mm; D/V 8.5 mm; and (2) A/P -0.2 - -0.3 mm; M/L ± 2 mm; D/V 9 mm for a bregma-lambda distance of 9mm. 200-300 nL of virus were delivered into the targeted area via a micropipette. Animals that underwent surgery were allowed to recover in their home cages for at least a week before electrode implant or slice physiology experiments; at least 2 weeks was allowed after virus injections in BF before CNO administration and data collection to ensure that DREADDs reached the axonal terminals in V1m. For the short hairpin M1 ACh receptor knockdown experiments, virus injections were performed at least 2 weeks before slice electrophysiology experiments.

### Electrode Implants and Monocular derivation

Rats were implanted with electrode arrays as previously described (Hengen et al., 2013; Hengen et a., 2016; Torrado-Pacheco et al., 2021). Briefly, custom 16-channel tungsten wire (33 μm tip diameter, Tucker-Davis Technologies, TDT) arrays were implanted in V1m, secured using dental cement, and grounded to screws inserted in the skull above the frontal cortex. Total headcap weight was approximately 2 g. Finally, two braided steel wires were implanted deep in the nuchal muscle for EMG recordings. MD was performed as described (Torrado Pacheco et al., 2021), except that eyelids were trimmed before suturing: rats were anesthetized with isoflurane, eyelids were trimmed and sutured (4 mattress sutures), and sutures were checked daily to ensure they remained intact.

### Continuous single-unit recordings in freely behaving animals and automated spike extraction, clustering, and sorting

Continuous single-unit recordings, spike extraction, clustering, and sorting were performed using our previously described pipeline (Hengen et al., 2016; Torrado Pacheco et al., 2021). After two full days of recovery, siblings were transferred to a clear plexiglass recording chamber for an additional day of habituation prior to the start of recordings, where they were separated into two arenas by a clear plastic divider with 1” holes to allow for tactile and olfactory interaction. Spikes were then recorded continuously for 7 days (except for a brief period during MD); for spike extraction, spikes were detected in the raw recorded signal as threshold crossings (-4 standard deviations from mean signal) and re-sampled at 3x the original rate. Custom scripts in Matlab and Python were used to cluster spikes from each channel for the entire recording period, and were classified as noise, multi-unit, or single unit. Only single-unit clusters with a clear refractory period were used for firing rate analysis. We classified units as RSU or FS based on established criteria (mean waveform trough-to-peak and tail slope). Only RSUs (putative excitatory neurons) were used for analysis. To establish “on” and “off” times for neurons, we used ISI contamination: when hourly % of ISIs > 3 ms was above 4%, units were considered offline. Based on these “on” and “off” times, only units that were online for at least 70% of the full 7-day experiment were used for analysis.

### Semi-automated behavioral state scoring

Behavioral state scoring was performed as previously described (Hengen et al., 2016; Torrado-Pacheco et al., 2021), except animal movement was tracked from video recordings using DeepLabCut (Mathis et al., 2018). Briefly, local field potentials (LFPs), EMG signals and animal movement were used for behavioral classification. For behavior classification, we used an in-house custom graphical user interface (GUI) built in MATLAB for semi-automated detection of 4 states: NREM sleep (high delta power, low theta power; low EMG and movement); REM sleep (low delta power; high theta power; lowest EMG and movement); quiet wake (low delta and theta; low EMG and movement); active wake (low delta and theta; higher EMG and movement).

### Behavioral State Analysis

The behavioral analysis shown in Figure 1F and in Table 1 was performed on the four ‘Hm4di + CNO’ animals in Figure 2, plus two more animals that had the same virus injection and array implant procedures, but they did not undergo MD (77.5% and 65.86% of ChAT+ neurons in the HDB were also RFP+ in these animals). All 6 animals for this analysis received 4 subcutaneous CNO injections total, spaced 12 hours apart, around ZT 00:00 and ZT 12:00 on two consecutive days. Once behavioral state data was scored for each animal, percent time spent in each of four behavioral states was calculated for each animal in the 8 hours immediately following each CNO injection, and in an equal number of 8 hour circadian-matched baseline periods outside of those CNO injection periods. Each CNO period was then normalized to the average of the two circadian-matched baseline periods. Since each animal received 4 CNO injections per experiment, the resulting 4 normalized values were averaged for each animal and then plotted, so that each data point in Figure 1 represents an animal.

**Table 1:** Animal behavior with and without HDB ACh inhibition.

| Time Period | REM (%) | NREM (%) | AW (%) | QW (%) |
| --- | --- | --- | --- | --- |
| Baseline (Light Periods) | 14.3±1.0 | 54.5±3.2 | 5.7±1.0 | 25.5±2.3 |
| CNO (Light Periods) | 11.9±1.3 | 55.9±3.8 | 6.2±0.7 | 25.9±2.5 |
| Baseline (Dark Periods) | 10.7±1.2 | 36.1±4.7 | 10.4±2.5 | 42.8±3.7 |
| CNO (Dark Periods) | 8.7±1.5 | 36.3±3.1 | 11.0±25.7 | 44.0±2.6 |

Percent time animals spent in each state during an 8-hour period in each corresponding condition, expressed as mean±SEM. N = 6 ChAT-Cre rats, all of which had Hm4di in the HDB.

**Figure 2:**
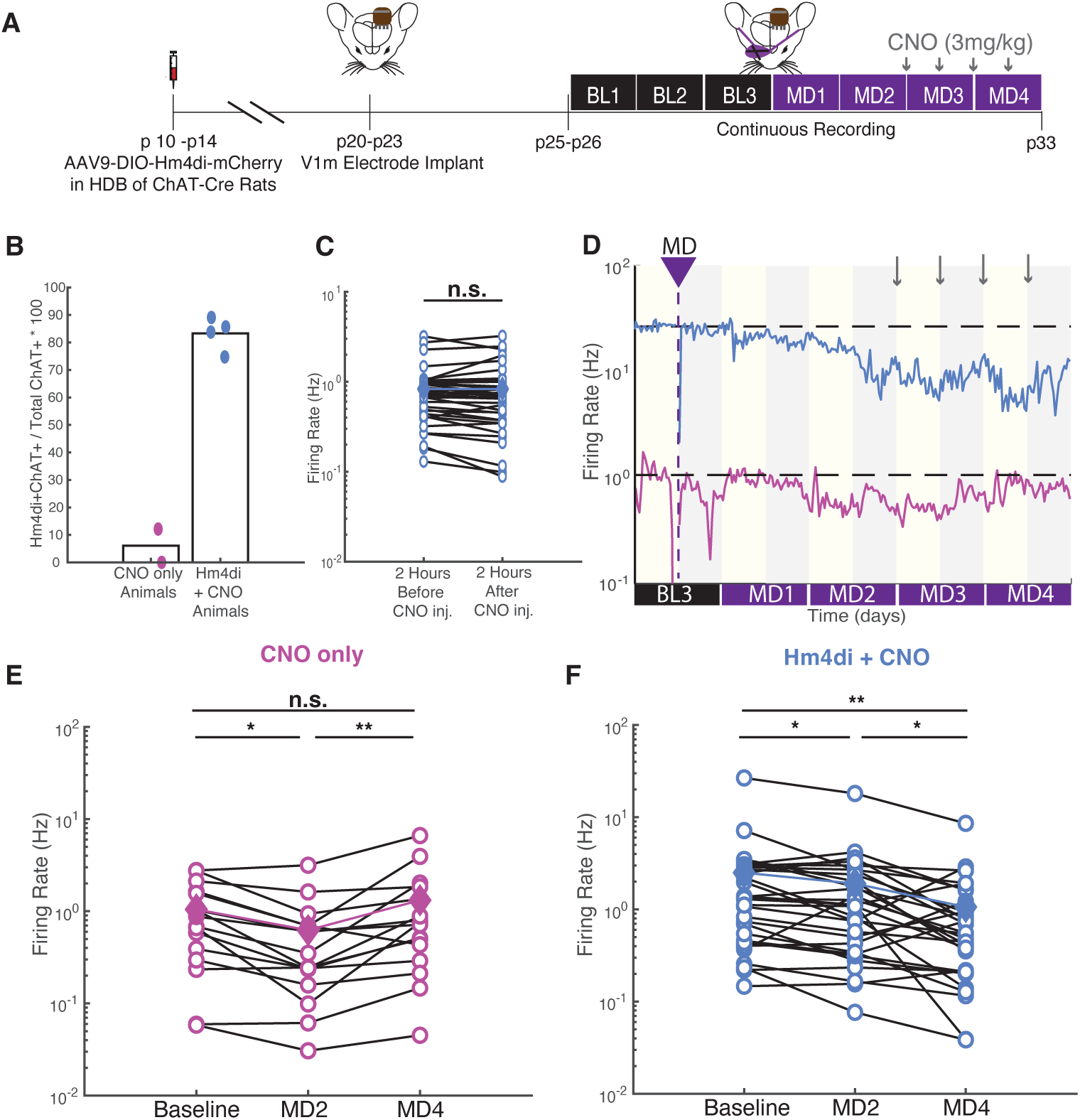
Cholinergic inhibition prevents upward firing rate homeostasis. A) Experimental timeline. All but one animal underwent virus injection surgery at or before p14. After at least one week of recovery, animals underwent electrode array implant in monocular visual cortex. After at least 3 more days of recovery, continuous recording began and lasted for 7 days. All experiments were complete before p33, and thus occurred within the classical visual system critical period. During the dark period on the third day of baseline, monocular deprivation (MD) was performed on the eye contralateral to the electrode implant. During the third and fourth days of MD, animals received 4 total subcutaneous CNO injections, one every 12 hours starting at ZT0 on MD3. B) Quantification of virus expression in ChAT+ cells in the horizontal diagonal band (HDB) of all animals used in these *in vivo* experiments. One animal that underwent virus injection showed very little virus expression (12% ChAT+ neurons in HDB were also mCherry+), so we included that animal in the “CNO only” cohort. The “CNO only” cohort also included another animal that did not receive any virus injection. In contrast, the four animals that were included in the “Hm4di + CNO” cohort had 75%, 88%, 85%, and 83% of ChAT+ neurons in the HDB also mCherry+. C) Inhibition of HDB ACh neurons does not acutely change firing rates of V1m regular spiking units. The firing rates of neurons from animals in the “Hm4di + CNO” cohort was indistinguishable in the 2 hours immediately preceding each CNO injection compared to the 2 hours immediately after each CNO injection. Empty circles each represent one cell, solid diamonds and colored line represent the mean firing rate at each time point. Mean FR before = 0.83 Hz, Mean FR after = 0.83 Hz. Median FR before = 0.70 Hz, Median FR after = 0.68 Hz. N = 37 cells from 4 animals. Wilcoxon sign rank test; before vs. after: p = 0.36. D) Firing rate traces of two example deprived cells over the course of the monocular deprivation, one from the “Hm4di + CNO” cohort, and one from a “CNO only” animal. E) and F) Ladder plot showing the activity of each deprived cell in the “CNO only” (**E**) and “Hm4di + CNO” (**F**) cohort averaged over a 12-hour period on baseline day 3, MD2, and MD4 in the open circles. The solid diamonds show the mean firing rate of all cells at each time point. The activity of deprived cells in **E** is significantly lower than baseline on MD2, but back up to baseline levels on MD4, while the activity of deprived cells in **F** is significantly lower than baseline on MD2 and still significantly lower than baseline on MD4. N = 16 cells from 2 animals (**E**) and 30 cells from 4 animals (**F**). Wilcoxon sign rank test with Bonferroni correction for multiple comparisons. **E**: Baseline vs. MD2: p = 0.011; MD2 vs. MD4: p = 0.007; Baseline vs. MD4: p = 1. **F**: Baseline vs. MD2: p = 0.040; MD2 vs. MD4: p = 0.010; Baseline vs. MD4: p = 0.003.

### LFP Band Analysis

The same animals and time periods were used here as in the *behavioral state analysis* above. LFP was extracted as previously described (Hengen et al., 2016; Torrado Pacheco et al., 2021), and the power in each frequency band (delta 0.5-4 Hz; theta 6-8 Hz; alpha 9-15 Hz) during each CNO period was normalized to the corresponding baseline periods. The resulting 4 normalized values per animal were averaged to give 1 data point per animal.

### In Vivo Firing Rate Analyses

To obtain firing rate estimates for individual RSUs from spike timestamps we computed spike counts in 30-minute bins throughout the experiment. To calculate mean firing rates in 12-hour periods, we took the average firing rate across all bins in that period. Baseline firing rates were calculated from the 12 hours right before MD, early MD firing rates were calculated from the first 12 hours on MD day 2, and late MD firing rates were calculated from the last 12 hours on MD day 4. These firing rate estimates and Baseline, MD2, and MD4 calculations are used throughout Figure 2.

### Drug Administration

For subcutaneous injections, water soluble CNO (Hello Bio) was dissolved in 0.9% sterile saline to reach the desired concentration (1 mg/ml). For drinking water CNO administration, CNO was dissolved in water to reach the desired concentration (0.05 mg/ml), and 10 mM saccharine chloride was added to the solution before giving to the animals. For DREADD (Hm4di) activation, CNO was administered via subcutaneous injections at a dose of 3 mg/kg every 12 hours, or via drinking water *ad libitum*. When administered via drinking water, animals drank about 15-20 mL per day to reach a dose of about 10-15 mg/kg over the course of 24 hours.

### Ex vivo brain slice preparation and Electrophysiology

*Ex vivo* slice preparation and electrophysiology was performed as previously described (Wen and Turrigiano, 2021), except that rats were used instead of mice, and V1m was identified by comparing the white matter morphology of the slice to those from the coronal sections illustrated in Paxinos and Watson (1998). Briefly, 300 μm thick slices from both hemispheres of V1 were obtained from rats and, after ∼45 minutes recovery, slices were used for electrophysiology 1-6 hours after slicing. L2/3 pyramidal neurons were identified based on slice and cell shape morphology and confirmed post-hoc with biocytin fill reconstructions. Borosilicate glass pipettes were pulled to achieve pipette resistances between 4 and 8 MΩ and were filled with K+ gluconate-based internal solution. Data were acquired using the open-source MATLAB-based software WaveSurfer (Janelia Research Campus, Howard Hughes Medical Institute), and data analysis was performed using in-house scripts written in MATLAB. For all analyses, neurons were excluded if access resistance was >20 MΩ, input resistance was <90 or >350 MΩ, Capacitance was < 50 pF, or membrane potential was greater than -55 mV

### mEPSC recordings

For spontaneous mEPSC recordings, slices were superfused with standard ACSF containing a drug cocktail of tetrodotoxin (TTX; 0.1 μM), aminophosphonopentanoic acid (AP-5; 50 μM), and picrotoxin (25 μM) to isolate AMPAR-mediated mEPSCs. L2/3 pyramidal neurons were targeted and held at -70 mV in whole-cell voltage clamp. Each neuron was recorded for 3-5 minutes in a series of 10 s sweeps, and a 500 ms 5 mV hyperpolarizing voltage step was administered at the beginning of each sweep so that passive properties could be monitored throughout the recording. Several 10 s traces were analyzed for each cell to ensure that 200-300 events were analyzed. If cells were not stable for at least 200 seconds of recording, or if the RMS exceeded 3.8 pA, the cell was excluded. Thereafter, to reliably detect mEPSC events and limit selection bias, in-house software with an automated template-based detection method was used. Event exclusion criteria included amplitudes less than 5 pA or rise times greater than 3 ms. Rise time (10-90%) and decay time constants (first order exponential fit) were calculated from the waveform average traces for each neuron.

### Intrinsic Excitability measurement

For intrinsic excitability measurements, slices were superfused with standard ACSF containing AP-5 (50 μM), picrotoxin (25 μM), and 6,7-dinitroquinoxaline-2,3-dione (DNQX; 25 μM) to block synaptic currents. L2/3 pyramidal neurons were held in current clamp with a small DC injection to maintain the resting membrane potential at -70 mV. Frequency-current (*f*-I) curves were obtained by delivering a series of 20500 ms current injections in amplitude increments of 20 pA. Passive properties were monitored in voltage clamp before and after current steps. In addition to passive property exclusion criteria, neurons were excluded if they displayed non-regular spiking firing patterns. Mean instantaneous firing rate (IFR) is defined as the mean reciprocal of the first two inter-spike intervals. Spike adaptation index is defined as the sum of inter-spike intervals during a 300 pA current step, where higher values indicate more adaptation.

### Carbachol Response experiments

To assess the functional efficacy of our M1 knockdown virus, slices were superfused with standard ACSF containing AP-5 (50 μM), picrotoxin (25 μM), and DNQX (25 μM). We recorded from control and M1 shRNA infected (identified by presence of YFP) neurons in current clamp with a small DC injection to maintain the resting membrane potential at -70 mV. After several minutes of baseline recording, we washed in ACSF with the same synaptic blockers as above, plus 10μM Carbachol (CCh) for 10 minutes, then washed out the CCh by superfusing back in the standard ACSF for 10 more minutes. Throughout the entire recording, every 5 seconds, we injected a hyperpolarizing 100 pA current injection for 500 ms, and once every 6 cycles of this, we instead injected a depolarizing 250 pA current injection for 500 ms to monitor passive properties and firing properties of the cells. For quantifications, we averaged values obtained during the first two minutes of baseline recording for baseline values, during the last two minutes of CCh wash-in for CCh values, and during the last two minutes of recording during CCh wash-out for wash values. IFR is defined as above, and after depolarization potential (ADP) is defined as the maximum membrane potential reached in the 1 second following the end of the depolarizing current injection.

### M1 shRNA generation

The shRNA sequence (5’-GAACCCACTGCATCTACATTT-3’) used for M1 knockdown was designed with the TRC algorithm (<u>Broad Institute</u>) and cloned into the pAAV-shRNA-ctrl vector from Addgene. pAAV-shRNA-ctrl was used as the empty vector plasmid in Figure 5. Our M1 shRNA plasmid was packaged in AAV9 viral vectors by Duke Viral Core. A modified version of the pAAV-shRNA-ctrl plasmid (eYFP was replaced with eGFP) had already been packaged in an AAV9 viral vector by a previous member of the lab, Lauren Tereshko, and this was used for empty vector *in vivo* viral injections.

#### Immunostaining

To recover cellular morphology after whole-cell recording, slices were post-fixed in 4% paraformaldehyde (PFA) for at least 24 hours, and then transferred to 0.01M PBS for storage before staining. For immunostaining experiments, animals were anesthetized with a ketamine/xylazine/acepromazine cocktail and transcardially perfused with 4% PFA. Brains were then removed and postfixed in PFA for at least 24-48 hours before sectioning V1 and the basal forebrain into 70 μm slices. Slices were washed in PBS before being incubated in a blocking solution (0.3% Triton X-100, 0.05% NaN_3_, 5% goat serum, and 3% BSA in 0.01 PBS) for 1 hour. Blocked slices were then incubated in the same blocking solution with primary antibodies added (ck antiGFP 1:350; gt antiChAT 1:100; rb antiRFP and ms antiRFP 1:500; rb anticFos 1:500; antiNeuN 1:500) at 4°C for 24 hours. The following day, slices were washed in PBS and then incubated in a solution (0.05% NaN_3_, 5% goat serum, and 3% BSA in 0.01 M PBS) containing secondary antibodies (streptavidin 1:350, all others 1:500). For immunostaining that involved the goat anti-ChAT primary antibody, goat serum was excluded from all solutions, and a sequential staining protocol, where slices were incubated with the anti-ChAT primary antibody overnight first, then washed and incubated again with any other primary antibodies overnight, was adopted for better results. For secondary antibody incubation, acute slices were incubated at 4°C overnight, all other slices were incubated at room temperature for 3 hours. Images were obtained using a confocal microscope (model LSM880, Zeiss). For immunostaining neuronal cultures, dishes were fixed at DIV10 with a 4% PFA and sucrose solution for 15 minutes. Dishes were then washed with PBS 3x5 minutes, then incubated in 100 uL/dish of blocking solution for 30 minutes. Dishes were then incubated with primary antibodies (rb antiM1 1:500, ck antiGFP 1:500) overnight at 4°C. The next day, they were washed again in PBS, and then incubated with secondary antibodies (1:500) for 1 hour at room temperature.

#### Immunohistochemistry quantification

##### cFos staining

To quantify cFos expression in Hm4di+ HDB ACh neurons, we created maximum projection images from z-stacks taken with a 20x objective in the HDB of fixed slices from Hm4di-injected animals. Images were background subtracted with a rolling ball radius of 50 pixels. RFP+ neurons were traced manually from the RFP channel, and the cFos+ neurons were identified by thresholding in the cFos channel of the image. Then all RFP labeled and RFP/cFos double labeled neurons were quantified from all HDB images from a given animal to find a total percentage per animal using the formula: (RFP+cFos+ / RFP+) * 100. As an additional no CNO control, we measured cFos expression in HDB ACh neurons with no viral Hm4di expression, using an anti-ChAT antibody rather than RFP. The quantification of cFos expression was performed in the same manner in these animals, except that neurons were identified by ChAT expression rather than RFP. We present the data and statistics with the two no CNO groups combined in Figure 1, but we note that cFos expression was significantly lower in Hm4di+ neurons after both 12 and 48 hours of CNO exposure compared to the Hm4di+ no CNO group alone as well.

##### Virus expression

To quantify how much Hm4di expression was present in each virus injected animal, we created maximum projection images from z-stacks taken with a 20x objective in the horizontal diagonal band of each animal. We then manually counted the number of cells that were ChAT labeled and RFP/ChAT double labeled and quantified the total percentage per animal using the formula: (ChAT+RFP+ / ChAT+) * 100.

##### M1 shRNA efficacy

To quantify M1 protein expression in transfected, cultured neurons, we created maximum projection images from z-stacks taken with an oil immersion 60x objective. Analyses were carried out over 4 dissections, and each dissection included several dishes each of neurons transfected with an empty vector or our M1 shRNA construct. Cells were only used if they were YFP+ to indicate successful transfection. In one dissection, we had no healthy M1 shRNA transfected neurons, so we only included the empty vector transfected neurons from that dissection in our analysis. Somas of each neuron were manually traced, and the mean intensity in the M1 channel was recorded for each cell. Each neuron’s M1 intensity was then normalized to the average empty vector neuron M1 intensity for that dissection, so that we could pool data across dissections and imaging sessions. To quantify M1 shRNA viral efficiency *in vivo*, we injected 4 rats bilaterally with our shRNA viral vector, sacrificed them two weeks later, and performed immunostaining on fixed slices. We then quantified the percent of YFP+ cells that were also NeuN+ using the formula: (YFP+NeuN+ / YFP+) * 100.

##### Statistical analyses

For each experiment, unless otherwise noted, results are reported as the mean ± SEM; sample sizes (both the number of neurons and the number of animals), statistical tests used, and *p* values are given either in the corresponding Results section or in the figure legends. For normally distributed data, a t-test was used for pairwise comparisons, and a one-way ANOVA followed by Tukey’s post hoc correction was used for multiple comparisons (n>2 groups). For non-normally distributed data, a Wilcoxon sign rank (unpaired data) test or a Wilcoxon rank sum (paired data) test was performed for pairwise comparisons; a Wilcoxon rank sum test with a Bonferroni post hoc correction was used for paired data with multiple comparisons, and a Kruskal-Wallis test followed by Tukey’s post hoc correction was used for multiple comparisons with unpaired data. For statistical tests that involved a Bonferroni correction for multiple comparisons, p values above 1 were replaced with 1. For distribution comparisons, a two-sample Kolmogorov-Smirnov test was used. Results were considered significant if p<0.05. Significance symbols used in figures are defined as such: *p<0.05; **p<0.01; ***p<0.001; n.s., not significant.

#### Data Availability

All data generated in this study are included in the manuscript and available upon request.

## RESULTS

### Long-lasting inhibition of V1-projecting HDB ACh neurons using DREADDs

We wished to know whether cholinergic neuromodulation of V1 is necessary to enable upward firing rate homeostasis and homeostatic plasticity. The major source of cholinergic projections to V1 arise from the horizontal diagonal band (HDB) subregion of the basal forebrain (BF). To chronically suppress the activity of these neurons, we used a viral vector expressing a Cre-dependent form of the inhibitory DREADD Hm4di (AAV9-hSyn-DIO-hM4D(Gi)-mCherry). By injecting this vector into the HDB of ChAT-Cre rats and waiting two weeks, we obtained robust expression in HDB ACh neurons (Fig. 1A,B; left panels), as well as in their axonal projections to V1 (Fig. 1A,B; right panels).

We next confirmed that Hm4di expression was able to effectively inhibit HDB ACh neurons upon administration of the exogenous ligand CNO. At least one week after viral delivery of Hm4di, ChAT-cre rats were given *ad libitum* access to CNO in their drinking water (Wen and Turrigiano, 2021) for either 0, 12, or 48 hours prior to sacrifice at ZT 01:00; animals were kept awake for 1 hour prior to sacrifice (ZT 00:00-01:00) using gentle handling to ensure some baseline level of HDB ACh neuron activity. We then performed immunostaining on HDB slices and probed for RFP expression (indicating an Hm4di+ cell) and cFos expression (an indicator of neural activity). We found that both 12 and 48 hours of CNO produced a ∼50% reduction in the fraction of HDB ACh neurons that were cFos+ (Figure 1C,D), indicating that we can achieve a long-lasting inhibition of their activity. As an additional control, we quantified cFos expression in HDB ACh neurons with no viral DREADD expression (using an anti-Chat antibody rather than RFP expression); these two control groups had very similar levels of baseline cFos expression (no virus, no CNO: 19.27±1.57%; Hm4di, no CNO: 21.53±3.72%), so these two control groups were combined (Figure 1D).

It has been shown that manipulation of BF ACh neurons can cause significant changes in cortical rhythms. Optogenetic activation causes reliable desynchronization of these rhythms, including decreases in delta band power and increases in theta band power (Pinto et al., 2014; Han et al., 2014; Xu et al., 2015), while inhibiting them causes a small increase in delta band power (Buzsaki et al., 1988; Pinto et al., 2014; Chen et al., 2016). To determine whether HDB inhibition affects brain-state activity within V1, we used local field potential (LFP) recordings from V1 to quantify the average LFP power in three different frequency bands (Delta, Theta, and Alpha) in the 8 hours immediately following subcutaneous CNO injections (3mg/kg), normalized to time- and circadian-matched periods from the same animals on recording days with no CNO administration. After CNO administration, when HDB ACh neurons were inhibited, delta power was significantly increased, while alpha power was significantly decreased (Figure 1E,F). This is consistent with the expected impact of BF inhibition on neocortical activity (Buzsaki et al., 1988; Pinto et al., 2014; Chen et al., 2016), indicating that inhibition of HDB ACh neurons produces measurable effects within V1.

In addition to altering cortical activity, acute activation of BF ACh neurons can cause sleep to wake transitions, while chronic activation can increase wakefulness (Han et al., 2014; Chen et al., 2016; Zant et al., 2016); in contrast, chronic inhibition of BF ACh neurons is reported to have only small and variable effects on sleep and wake states (Blanco-Centurion et al., 2006; Kaur et al., 2008; Fuller et al., 2010; Chen et al., 2016). To determine the impact of HDB ACh neuron inhibition, we used LFP, electromyography (EMG), and video recordings to quantify the amount of time Hm4di-injected animals spent in each of four behavioral states (Active Wake, AW; Quiet Wake, QW; Rapid Eye Movement sleep, REM; and Non-REM NREM sleep) in the 8 hours immediately following CNO administration, normalized to time- and circadian-matched periods with no CNO administration. We found that rats spent comparable amounts of time in AW, QW, and NREM sleep before and after HDB ACh inhibition, but there was a small decrease in time spent in REM sleep (Figure 1G, Table 1). Because rats normally spend little time in REM sleep (∼10%, see Table 1), this small reduction in REM corresponds to very little change in total sleep. Moreover, because upward firing rate homeostasis occurs almost exclusively during wake (Hengen et al. 2016), this small change in REM is unlikely by itself to impair upward firing rate homeostasis. Taken together, our results show that we can chronically reduce the activity of cholinergic projections to V1 without dramatically influencing the behavioral state of the animal.

### Cholinergic inhibition prevents upward firing rate homeostasis

We showed previously that 2 days of monocular deprivation (MD) first reduces firing rates in V1, but that over the next 2 days, firing rates are homeostatically regulated back to the pre-MD baseline (Hengen et al., 2016; Torrado Pacheco et al., 2021; Tatavarty et al., 2020). To determine whether activity of HDB ACh neurons is necessary for this upward firing rate homeostasis (FRH), we obtained continuous extracellular single unit recordings from monocular visual cortex (V1m) of freely behaving animals during MD, and then inhibited HDB ACh neurons on MD 3 after firing rates had fallen (Figure 2A). First, we replicated our previous demonstrations of upward firing rate homeostasis in two ‘CNO only’ control animals, one with no viral injection into the HDB, and one where the injection failed and DREADDs expression was minimal (12% of ChAT+ neurons in HDB were RFP+, vs. 88%, 85%, 83%, and 75% in animals in the ‘Hm4di+CNO’ group; Figure 2B). We found that MD first depressed firing, but over the next two days, firing returned to baseline despite continued MD, as expected (Fig. 2D,E; Hengen et al., 2016; Torrado Pacheco et al., 2021). In contrast, inhibition of HDB ACh neurons during MD 3-4 completely prevented the normal recovery of firing rates, and instead firing rates on average fell even further (Fig 2D,F, n = 4 animals). Importantly, we found that HDB ACh inhibition had no acute effect on V1m firing rates (Figure 1C). Thus, upward FRH in V1m is dependent on HDB ACh activity.

### Cholinergic inhibition impairs synaptic scaling

Two major forms of homeostatic plasticity are known to contribute to upward firing rate homeostasis in L2/3 pyramidal neurons: synaptic scaling and intrinsic homeostatic plasticity (Lambo and Turrigiano, 2013; Hengen et al., 2016; Tatavarty et al., 2020). We thus hypothesized that inhibiting wake-active HDB inputs to V1 prevents upward FRH by disrupting one or both of these forms of homeostatic plasticity.

To test this hypothesis, we used a simplified paradigm to directly suppress the activity of L2/3 pyramidal neurons in V1m using inhibitory DREADDs (Fig. 3A); this paradigm has been shown to induce both synaptic scaling up and intrinsic homeostatic plasticity in L2/3 pyramidal neurons of young mice (Wen and Turrigiano, 2021). We virally delivered Hm4di under the control of a CaMKII promoter (AAV9-CaMKII-hM4D(Gi)-mCherry) to one hemisphere of V1m, rats were given *ad libitum* access to CNO drinking water (0.05 mg/mL) for 24-48 hours prior to slice electrophysiology (Figure 3A,B), and L2/3 pyramidal neurons were then targeted for whole-cell recordings (Fig. 3B, C). We first confirmed that activity suppression with Hm4di + CNO in L2/3 V1m pyramidal neurons indeed induced synaptic scaling: Hm4di^+^ neurons had significantly larger miniature excitatory post-synaptic currents (mEPSCs) than Hm4di^-^ neurons after 48 hours of CNO treatment (CNO only, 10.6±0.1 pA vs. V1 Hm4di + CNO, 11.2±0.1). In contrast, when the HDB was inhibited at the same time as V1 (by expressing Cre-dependent Hm4di in HDB ACh neurons), we no longer saw a significant increase in mEPSC amplitude (Fig. 3D; HDB ACh Hm4di + CNO, 11.0±0.1 vs. V1 + HDB ACh Hm4di + CNO, 11.4±0.2). In comparing all four conditions, however, we noticed that both conditions with HDB ACh inhibition had slightly larger amplitude mEPSCs than the CNO only condition (Fig. 3D, V1 + HDB ACh Hm4di + CNO significantly different from CNO only). There was no significant difference in mEPSC frequency in any group (Figure 3E), and only subtle differences in passive properties (Table 2). These data suggest that HDB ACh inhibition alone causes a small increase in mEPSC amplitude in L2/3 pyramidal neurons and also inhibits or possibly occludes the induction of synaptic scaling. These results suggest that ACh signaling in V1 is important for the normal induction of synaptic scaling, but they do not provide a clear mechanism by which HDB ACh inhibition prevents firing rate homeostasis during prolonged MD.

**Figure 3:**
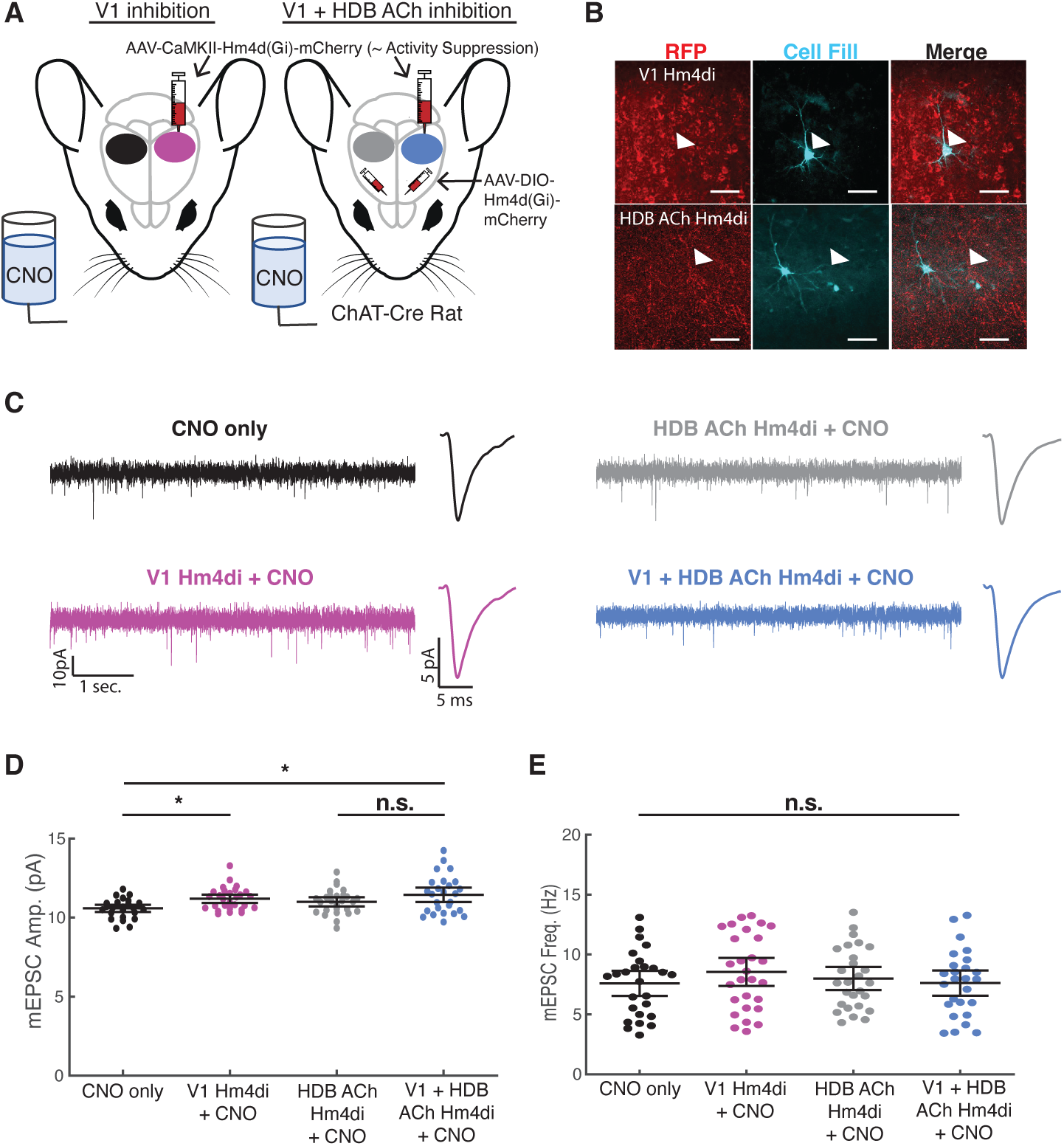
Cholinergic inhibition impairs synaptic scaling. A) Schematic of experimental design. Virus expressing Hm4di driven by a CaMKII promoter was injected unilaterally in V1m of each rat, with the other hemisphere serving as a within-animal control. In one set of ChAT-cre rats, virus expressing Cre-dependent Hm4di was also injected bilaterally in the HDB. All animals had access to *ad libitum* water with CNO for 48 hours prior to sacrifice for *ex vivo* electrophysiology. B) Example images of Hm4di expression and biocytin cell fill from recorded V1m L2/3 pyramidal cells. Top: Hm4di expression (RFP) driven by CaMKII in visual cortex; white arrowhead indicates the filled Hm4di+ neuron. Bottom: white arrowhead indicates Hm4di+ projections (RFP) in V1m near a recorded cell; image taken from control hemisphere of an animal that received Cre-dependent Hm4di injections into the HDB. Scale bar = 50 μm. C) Representative mEPSC traces from each experimental group. D) and E) mEPSC amplitudes (**D**) and frequencies (**E**) of all recorded cells in each experimental group. Black lines indicate mean ± SEM. CNO only: 26 cells from 9 animals; V1 Hm4di + CNO: 28 cells from 9 animals; HDB ACh Hm4di + CNO: 27 cells from 6 animals; V1 Hm4di + HDB ACh Hm4di + CNO: 26 cells from 6 animals. **D**: Kruskal-Wallis test p = 0.017 with Tukey-Kramer post hoc. CNO only vs. V1 Hm4di + CNO: p = 0.041; CNO only vs. HDB ACh Hm4di + CNO: p = 0.29; CNO only vs. V1 + HDB ACh Hm4di + CNO: p = 0.021; V1 Hm4di + CNO vs. HDB ACh Hm4di + CNO: p = 0.82; V1 Hm4di + CNO vs. V1 + HDB ACh Hm4di + CNO: p = 0.99; HDB ACh Hm4di + CNO vs. V1 + HDB ACh Hm4di + CNO: p = 0.671. **E**: Kruskal-Wallis test p = 0.67.

**Table 2:** Passive neuronal properties and kinetics for mEPSC experiments in Figure 3.

|  | CNO only | V1 Hm4di + CNO | HDB ACh Hm4di + CNO | V1 + HDB ACh Hm4di + CNO | P value |
| --- | --- | --- | --- | --- | --- |
| **R<sub>in</sub> (MΩ)** | 161.9±9.8 | 151.4±7.5 | 163.0±7.8 | 175.3±9.5 | 0.29 |
| **V<sub>rest</sub> (mV)** | -76.5±1.0 | -77.9±1.0 | -75.5±1.0 | -77.3±0.8 | 0.23 |
| **Cp (pF)** | 112.5±4.2 | 118.9±2.7 | 118.7±4.0 | 123.7±4.5 | 0.33 |
| **Rise Time**<br>**(ms)<sup>a</sup>** | 1.6±0.02 | 1.6±0.02 | 1.7±0.02 | 1.7±0.03 | 0.00005 |
| **Decay $\tau$ (ms)** | 3.0±0.16 | 2.6±0.09 | 3.0±0.13 | 3.0±0.18 | 0.09 |

Kruskal-Wallis test with Tukey-Kramer post hoc
<sup>a</sup> p = 0.00005. Pairwise comparisons: CNO only vs. HDB ACh Hm4di + CNO p = 0.013; V1 Hm4di + CNO vs. HDB ACh Hm4di p = 0.0002; V1 Hm4di + CNO vs. V1 + HDB ACh Hm4di + CNO p = 0.004

### Cholinergic inputs are necessary to maintain the intrinsic excitability of L2/3 pyramidal neurons during activity suppression

Intrinsic excitability is another major determinant of V1 firing rates (Maffei and Turrigiano, 2008; Lambo and Turrigiano, 2013; Trojanowski et al., 2021), so we next examined how HDB ACh inhibition affects intrinsic excitability. We used the same dual inhibition of V1 and HDB paradigm as above (Fig. 4A), except we recorded from L2/3 pyramidal neurons in current clamp in the presence of synaptic blockers and constructed FI curves by injecting a series of DC current steps to elicit spikes (Fig. 4B,C). DREADDs-mediated inhibition of V1m alone caused a small but not statistically significant increase in intrinsic excitability in L2/3 pyramidal neurons (Figure 4B-E “V1 Empty Vector + CNO” vs. “V1 Hm4di + CNO” conditions). Because the homeostatic regulation of intrinsic excitability is tightly developmentally regulated in these neurons (Wen and Turrigiano, 2021), these results suggest that after postnatal day (p)24 in rats, upward intrinsic homeostatic plasticity is significantly damped in these neurons. We then measured intrinsic excitability after inhibition of HDB ACh neurons alone, and again observed no significant change (Fig. B,C, “HDB ACh Hm4di + CNO” condition). However, much to our surprise, inhibiting HDB ACh neurons while simultaneously suppressing V1m activity caused a dramatic decrease in intrinsic excitability, characterized by a rightward and downward shift in the FI curve (Fig. 4C, “V1 + HDB ACh Hm4di + CNO” condition), and a significant decrease in the area under the FI curve (Fig 4D). This decrease in excitability was accompanied by a significant increase in rheobase (Figure 4E), with no change in adaptation index, input resistance, or other passive properties (Figure 4F,G, Table 3). This result suggests that cholinergic inputs to V1m are critical for maintaining the intrinsic excitability of L2/3 pyramidal neurons during activity suppression, and it is consistent with the ability of HDB ACh inhibition to prevent upward FRH (Figure 2).

**Figure 4:**
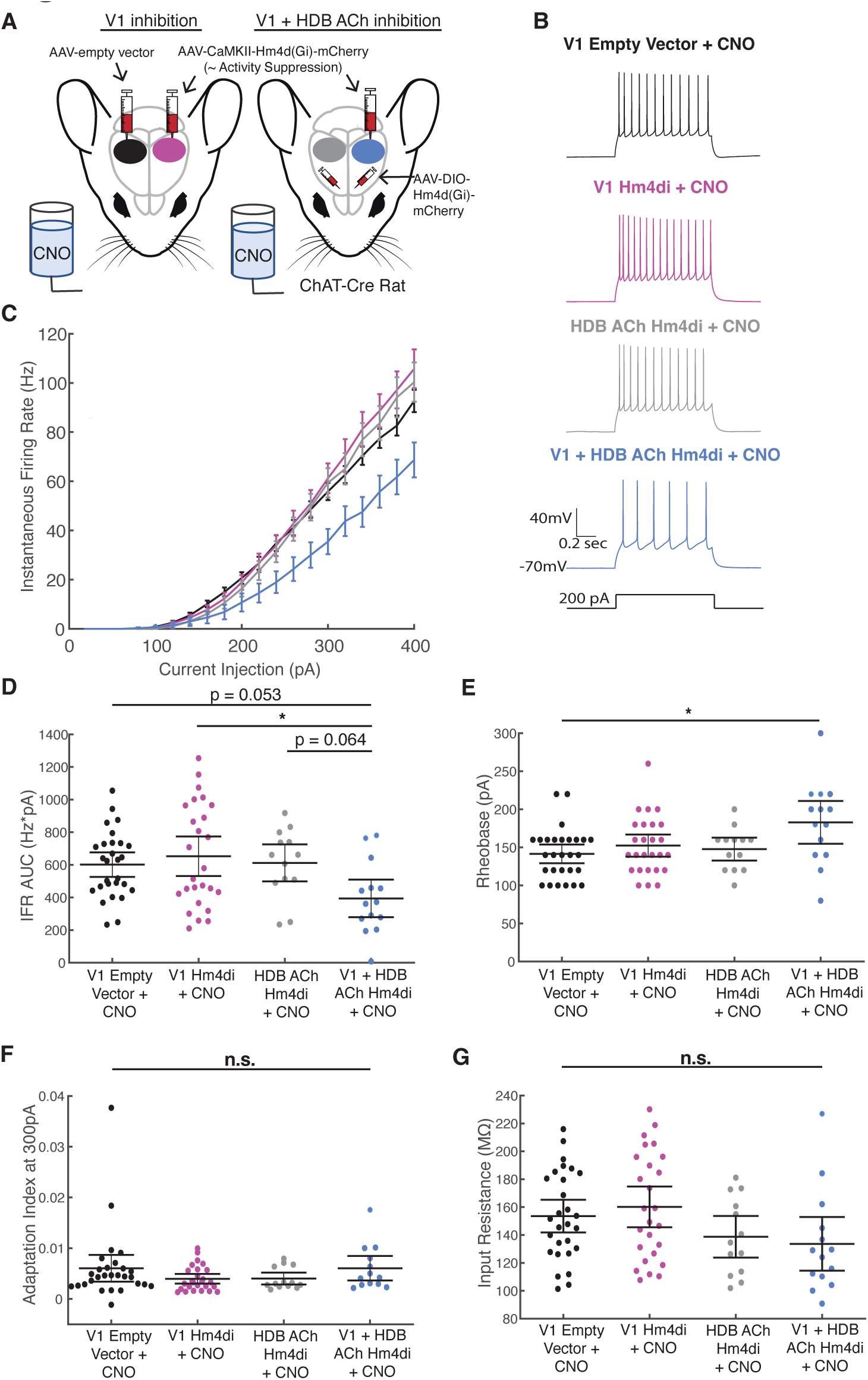
Cholinergic inputs are necessary to maintain the intrinsic excitability of L2/3 pyramidal neurons during activity suppression. A) Schematic of experimental design. All animals had access to *ad libitum* water with CNO for 24 hours prior to sacrificing for *ex vivo* electrophysiology. B) Representative recordings from V1m L2/3 pyramidal neurons from each experimental group evoked by 200 pA current injection. C) Firing rate-current injection (f-I) curves for each experimental condition. The y axis indicates mean instantaneous firing rate. D) Quantification of the area under each f-I curve, calculated individually for each neuron. Black lines indicate mean ± SEM. Kruskal-Wallis test p = 0.025 with Tukey-Kramer post hoc. V1 empty vector + CNO vs. V1 Hm4di + CNO: p = 0.98; V1 empty vector vs. HDB ACh Hm4di + CNO = 0.98; V1 empty vector + CNO vs. V1 + HDB ACh Hm4di + CNO = 0.053; V1 Hm4di + CNO vs. HDB ACh Hm4di + CNO = 1; V1 Hm4di + CNO vs. V1 + HDB ACh Hm4di + CNO: p = 0.027; HDB ACh Hm4di + CNO vs. V1 + HDB ACh Hm4di + CNO: p = 0.064. E) Rheobase for each recorded cell. Black lines indicate mean ± SEM. Kruskal-Wallis test p = 0.039 with Tukey-Kramer post hoc. V1 empty vector + CNO vs. V1 Hm4di + CNO: p = 0.75; V1 empty vector vs. HDB ACh Hm4di + CNO = 0.92; V1 empty vector + CNO vs. V1 + HDB ACh Hm4di + CNO = 0.021; V1 Hm4di + CNO vs. HDB ACh Hm4di + CNO = 1; V1 Hm4di + CNO vs. V1 + HDB ACh Hm4di + CNO: p = 0.18; HDB ACh Hm4di + CNO vs. V1 + HDB ACh Hm4di + CNO: p = 0.24. F) Adaptation index for each recorded cell, measured from recorded traces at 300 pA injected current. Kruskal-Wallis test: p = 0.29. G) Input resistance for each recorded cell. Kruskal-Wallis test: p = 0.063. Sample size for all plots: V1 Empty Vector + CNO: n = 28 cells from 7 animals; V1 Hm4di + CNO: n = 26 cells from 7 animals; HDB ACh Hm4di + CNO: n = 13 cells from 4 animals; V1 Hm4di + HDB ACh Hm4di + CNO: n = 14 cells from 4 animals.

**Table 3:** Passive neuronal properties and kinetics for intrinsic excitability experiments in Figure 4.

|  | **V1 Empty**<br>**Vector + CNO** | **V1 Hm4di +**<br>**CNO** | **HDB ACh**<br>**Hm4di + CNO** | **V1 + HDB ACh**<br>**Hm4di + CNO** | **P value** |
| --- | --- | --- | --- | --- | --- |
| **R<sub>in</sub> (MΩ)** | 153.6±5.7 | 160.2±7.3 | 138.7±7.1 | 133.7±8.9 | 0.063 |
| **V<sub>rest</sub> (mV)** | -79.1±0.8 | -78.3±0.7 | -74.8±1.1 | -76.7±1.2 | 0.055 |
| **Cp (pF)** | 118.0±2.3 | 118.1±3.2 | 126.7±3.0 | 126.7±3.9 | 0.104 |

Kruskal-Wallis test

### Knockdown of M1 ACh receptors *in vitro* and *in vivo*

Our results so far indicate that HDB ACh inputs to V1 are important for enabling upward FRH, and for maintaining the intrinsic excitability of L2/3 pyramidal neurons when their activity is suppressed. This raises the question of where in the brain ACh is acting to exert these effects. Inhibition of HDB ACh neurons could work through local release directly onto L2/3 pyramidal neurons, by targeting other cell types within V1, or by targeting other structures that indirectly affect activity or modulatory signaling within V1. To begin to tease apart these possibilities, we targeted the M1 ACh receptor, a Gq coupled metabotropic receptor that is highly expressed in rodent visual cortex, especially on somas and dendrites of pyramidal neurons, including in L2/3 (Levey et al., 1991; Yamasaki et al., 2010; Groleau et al., 2015).

We generated a short hairpin plasmid targeting the 3’UTR of the rat M1 ACh receptor, also expressing yellow fluorescent protein (YFP) for visualization. When this plasmid was transfected into cultured cortical neurons, we found a significant reduction in M1 AChR protein compared to neurons transfected with an empty vector, as assessed by immunohistochemistry (Figure 5A-C). We then packaged the shRNA into an AAV9 viral vector, targeted V1m and waited at least two weeks for full expression. We observed robust expression of our shRNA in multiple cell types in V1m, with about 62±7% of YFP+ cells expressing the pyramidal neuron marker NeuN (Figure 5D). This indicates our shRNA likely infected both pyramidal neurons and interneurons, but we restricted our recordings in subsequent experiments to YFP+ pyramidal neurons. To assess the functional efficacy of our M1 knockdown we obtained whole-cell current clamp recordings and activated ACh receptors by washing in carbachol (CCh), a stable ACh analog, while injecting a series of small depolarizing and hyperpolarizing current steps to measure changes in firing and passive properties (Figure 5E). Consistent with previous reports (McCormick and Prince, 1985; Gulledge and Stuart, 2005; Gulledge et al., 2009), CCh wash-in increased firing rate and the magnitude of the after-depolarization potential (ADP) in unmanipulated pyramidal neurons, and these effects reversed upon CCh wash-out. M1 knockdown completely abolished both of these effects (Figure 5F,G); while there was a small decrease in firing in the presence of CCh in these knockdown neurons, this effect was more pronounced after CCh wash-out, suggesting it is a run-down effect of the long recording time rather than a specific CCh response in neurons with reduced M1 expression. Together, these data show that we can effectively knock down M1 ACh receptor expression in V1m, and that in L2/3 pyramidal neurons, this receptor mediates most of the postsynaptic response to CCh application.

**Figure 5:**
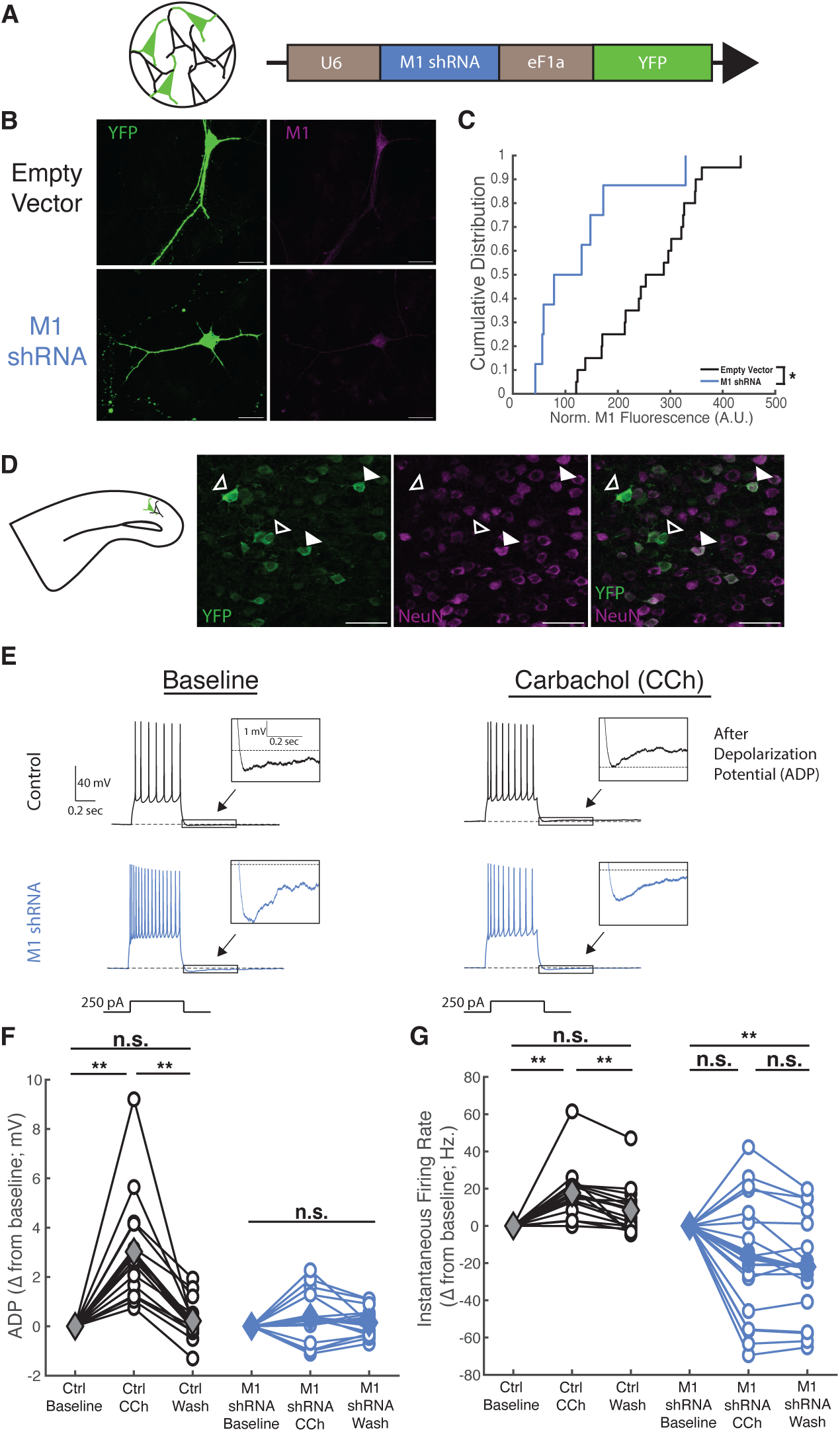
Knockdown of M1 ACh receptors *in vitro* and *in vivo*. A) Schematic indicating experimental system (neuronal cultures) and plasmid containing shRNA targeting M1 ACh receptors (M1 shRNA) used in B and C. B) Example images of cultured neurons transfected with an empty vector plasmid (top) or a plasmid containing M1 shRNA (bottom). Green = YFP; magenta = M1 ACh receptor. Scale bar = 20 μm. C) Cumulative distribution curves of M1 fluorescence intensity in somas of cultured neurons transfected with an empty vector (black) or M1 shRNA (blue); the M1 shRNA curve is significantly left shifted, indicating a reduction in M1 fluorescence intensity compared to empty vector. Cultures were fixed and stained after 3 days of transfection. Fluorescence intensities for each cell were normalized to the average fluorescence intensity of empty vector neurons in the corresponding group of cultures to account for any dissection related differences. Kolmogorov-Smirnov test p = 0.012. Empty vector: n = 20 cells from 4 dissections, m1 shRNA: n = 8 cells from 3 dissections. D) **Left**: schematic indicating *ex vivo* slice preparation for **D-G**. **Right**: Example image of an AAV9-M1 shRNA-YFP injection in V1m L2/3. 62±7% of YFP+ neurons were also NeuN+. Open arrow heads denote YFP+ cells, indicating expression of our shRNA, that did not express NeuN. Closed arrow heads denote YFP+ cells that also expressed NeuN. Green = YFP; magenta = NeuN. Scale bar = 50 μm. E) Control or knockdown neurons were recorded for several minutes of baseline in regular ACSF with synaptic blockers, followed by 10 minutes of Carbachol (CCh) wash-in and 10 more minutes of CCh wash-out. Representative traces from a control and a knockdown neuron in response to 250 pA of injected current, under baseline conditions (left) and in the presence of CCh (Right). Note that the control cell shows an increased firing rate and presence of an after-depolarization potential (ADP) in CCh compared to baseline, while the M1 shRNA cell does not show ADP in the presence of CCh and shows a slight decrease in firing rate, likely due to the long recording time. F) and G) Quantification of ADP (**F**) and instantaneous firing rate (**G**) for each recorded neuron at baseline, at the end of the CCh wash-in, and at the end of the CCh wash-out (open circles). The solid diamonds indicate the mean. All values are represented as a change from baseline. Control cells: n = 16 cells from 4 animals. M1 shRNA cells: n = 20 cells from 4 animals. Wilcoxon sign rank test with Bonferroni correction. **F**) Ctrl baseline vs. Ctrl CCh: p = 0.0013; Ctrl CCh vs. Ctrl Wash: p = 0.0013; Ctrl baseline vs. Ctrl Wash: p = 1; M1 baseline vs. M1 CCh: 0.17; M1 CCh vs. M1 wash: 1; M1 baseline vs. M1 wash: 0.84. **G**) Ctrl baseline vs. Ctrl CCh: p = 0.0016; Ctrl CCh vs. Ctrl wash: 0.0013; Ctrl baseline vs. Ctrl wash: p = 0.054; M1 baseline vs. M1 CCh: p = 0.12; M1 CCh vs. M1 wash: p = 0.17; M1 baseline vs. M1 wash: p = 0.0045.

### Maintenance of intrinsic excitability after V1 activity suppression is not mediated through V1 M1 ACh receptors

Having established that we can effectively knock down expression of M1 AChRs in V1m, we next asked whether this receptor mediates the activity-dependent maintenance of intrinsic excitability by cholinergic inputs to V1. To test this, we virally expressed either an empty vector, our M1 knockdown construct, or the combination of our M1 knockdown construct and CaMKII-Hm4di in V1m (Figure 6A). All animals then received CNO in their drinking water for 24 hours prior to slice recordings from V1m L2/3 pyramidal neurons to generate FI curves. Interestingly, there was no difference in intrinsic excitability between any of the conditions, with the FI curves completely superimposed (Figure 6B-E), and only subtle differences in passive properties (Table 4). Thus, even though M1 is the most abundant ACh receptor in rodent V1, reducing signaling through this receptor is not sufficient to mimic the effects of HDB inhibition. This suggests either that a less abundant ACh receptor type within V1 is the critical target, or that the effects of HDB inhibition on V1 plasticity are mediated indirectly via a target outside of V1.

**Figure 6:**
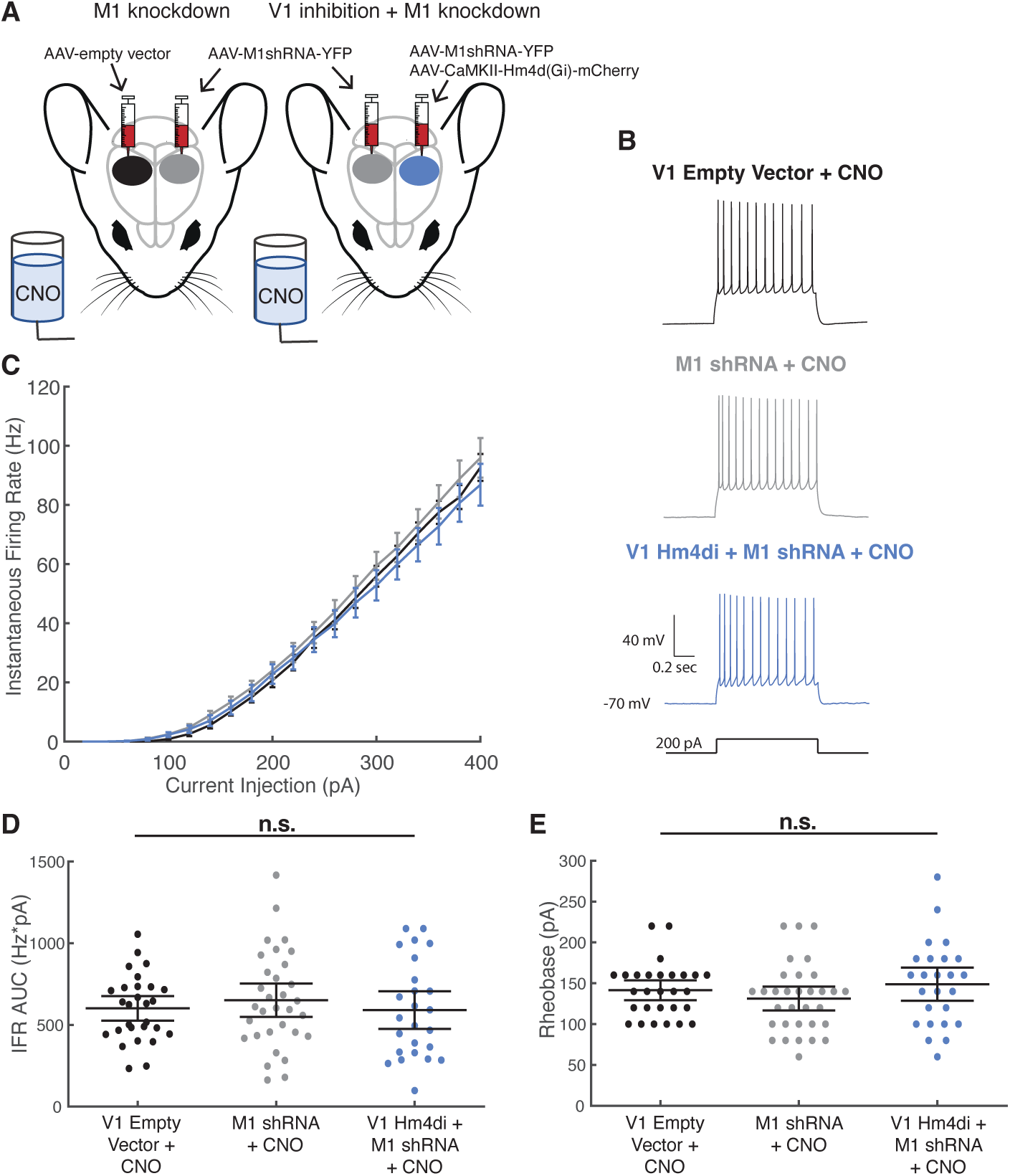
Maintenance of intrinsic excitability after V1 activity suppression is not mediated through V1 M1 ACh receptors. A) Schematic of experimental design. All animals had access to *ad libitum* water with CNO for 24 hours prior to sacrificing for *ex vivo* electrophysiology. B) Representative recordings from V1m L2/3 pyramidal neurons from each experimental group evoked by 200 pA current injection. C) F-I curves for each experimental condition. The y axis indicates instantaneous firing rate. D) Quantification of the area under each f-I curve, calculated individually for each neuron. Black lines indicate mean ± SEM. Kruskal-Wallis test: p = 0.73. E) Rheobase for each recorded cell. Black lines indicate mean ± SEM. Kruskal-Wallis test: 0.26.

**Table 4:** Passive neuronal properties and kinetics for intrinsic excitability experiments in Figure 6.

**Figure 6**
|  | **V1 empty**<br>**Vector + CNO** | **M1 shRNA +**<br>**CNO** | **V1 Hm4di + M1**<br>**shRNA + CNO** | **P value** |
| --- | --- | --- | --- | --- |
| **R<sub>in</sub> (MΩ)** | 153.6±5.7 | 169.7±8.0 | 173.1±10.6 | 0.35 |
| **V<sub>rest</sub> (mV)** | -79.1±0.8 | -78.5±0.8 | -76.5±0.9 | 0.16 |
| **Cp (pF)<sup>b</sup>** | 118.0±2.3 | 118.3±3.1 | 106.4±2.6 | 0.005 |

Kruskal-Wallis test with Tukey-Kramer post hoc
<sup>b</sup> p = 0.005. pairwise comparisons: V1 empty vector + CNO vs. M1 shRNA + Hm4di + CNO p = 0.015; M1 shRNA + CNO vs. M1 shRNA + Hm4di + CNO p = 0.010.

## DISCUSSION

Behavioral states play vital roles in the regulation and coordination of cortical plasticity, including the induction of firing rate homeostasis (Wang et al., 2011; Lee and Dan, 2012; Abel et al., 2013; Tononi and Cirreli, 2014; Hengen et al., 2016; Frank, 2017; Torrado Pacheco et al., 2021). Here we explore why upward FRH is confined to AW states, by testing the hypothesis that high levels of cholinergic neuromodulation are necessary for its expression. We find that chemogenetic inhibition of HDB ACh neurons, the major cholinergic inputs to V1, completely prevents upward FRH during monocular deprivation. We then ask how HDB ACh inhibition affects cellular plasticity mechanisms known to contribute to FRH, and we find that inhibition of HDB ACh neurons prevents synaptic scaling up and dramatically decreases the intrinsic excitability of activity-deprived V1 neurons. These data show that ACh plays an important role in the induction of homeostatic plasticity and is crucial for the dynamic maintenance of intrinsic excitability of L2/3 pyramidal neurons. Finally, knockdown of the abundant M1 ACh receptors in V1 neurons failed to phenocopy HDB ACh inhibition, suggesting that these effects are mediated through other receptors or other brain regions, or possibly non-neuronal cell types within V1 (Takata et al., 2011, Maurer and Williams, 2017). Taken together, our data show that wake-active cholinergic activity enables upward firing rate homeostasis through combined effects on synaptic and intrinsic plasticity, and they suggest more broadly that neuromodulatory tone is a critical factor that segregates upward and downward homeostatic plasticity into distinct behavioral states.

The difference between AW (when upward firing rate homeostasis occurs) and QW (when it does not) in the neocortex largely consists of differences in neuromodulatory tone. Specifically, cholinergic and noradrenergic (NA) modulation are both much higher in AW and contribute to creating the highly desynchronized cortical activity typical of this state (Constantinople and Bruno, 2011; Lee and Dan, 2012). Two main considerations made ACh a promising starting point for exploring neuromodulatory control of upward FRH. First, the target-specificity of BF ACh sub-regions (Kim et al., 2016) allowed us to target HBD inputs to V1 with greater specificity than possible if (e.g.) we were manipulating NA neurons in the locus coeruleus (Kim et al., 2016). Second, there is an extensive literature on the effects of ACh modulation on other forms of experience-dependent plasticity in the neocortex that provided a foundation for thinking about possible mechanisms by which ACh could be modulating homeostatic plasticity (Bear and Singer 1986; Pinto et al., 2014; Kang et al., 2015; Maurer and Williams 2017). Research has shown that ACh modulation is important for ocular dominance plasticity (Bear and Singer, 1986), causes long-term enhancement of visual cortex responses to salient stimuli (Pinto et al., 2014; Kang et al., 2015), and facilitates some forms of long-term depression in neocortex (Kirkwood et al., 1999; Caruana et al., 2011). ACh modulation has diverse effects on target neurons due to its multiple receptor subtypes and modes of release, with the emergent view that ACh modulation in cortex selectively potentiates neuronal responses to salient information (Gulledge and Stuart, 2005; Yamasaki et al., 2010; Picciotto et al., 2012; Alitto and Dan, 2013). Our results show that HDB ACh activity is critical for the induction of upward FRH, but they leave open the possibility that other neuromodulators, such as NA, could also contribute in a cooperative manner. Such cooperativity could provide a mechanism by which specific constellations of cellular plasticity mechanisms are on or offline during distinct brain states.

Reducing HDB ACh modulation had complex effects on the induction of two cellular forms of plasticity known to contribute to upward firing rate homeostasis: synaptic scaling up, and intrinsic homeostatic plasticity (Lambo and Turrigiano, 2013; Hengen et al., 2016). While inhibiting HDB ACh activity did prevent scaling up of mEPSCs in L2/3 pyramidal neurons induced by direct inhibition of activity in V1, it also produced a small increase in baseline mEPSC amplitude, making it difficult to differentiate between a complete block or partial block/partial occlusion of synaptic scaling. Further, the increase in baseline mEPSC amplitude induced by HDB ACh inhibition should act to increase activity of L2/3 pyramidal neurons, while in contrast we observed that firing rates of regular spiking/putative pyramidal neurons (measured across neocortical layers) failed to undergo a homeostatic increase. It is possible that the effect of HDB ACh inhibition on baseline mEPSCs is cell-type specific, and not evident in pyramidal neurons outside of L2/3, so that the net impact on circuit excitability is small. Regardless, these data show both that ACh modulation is important for the normal induction of homeostatic synaptic scaling up, and that there are likely effects on other plasticity mechanisms that contribute to the lack of firing rate homeostasis, such as intrinsic excitability changes.

We were surprised to find that DREADDs-mediated activity suppression in V1m L2/3 pyramidal neurons in our juvenile rats did not significantly increase intrinsic excitability, even though our experiments were carried out well within the classical visual system critical period (p24-p32; Figure 4), when this same method induces a robust increase in intrinsic excitability in similarly aged mice (p24-29; Wen and Turrigiano, 2021), and prolonged MD induces a robust intrinsic excitability increase in L2/3 neurons in binocular V1 of rats (p24-27; Lambo and Turrigiano, 2013). We previously found that intrinsic homeostatic plasticity is strongly developmentally regulated in V1m and is completely absent in adult mice (p45-55). Thus, one explanation for the differences between studies is that this developmental downregulation occurs earlier in rats than in mice. Intriguingly, we found that while inhibiting HDB ACh neurons alone had no effect on baseline intrinsic excitability, it caused a dramatic decrease in intrinsic excitability in V1m L2/3 pyramidal neurons when it was coincident with activity suppression in V1 (Figure 4). Thus, the combination of activity suppression and a lack of ACh input induces an anti-homeostatic regulation of intrinsic excitability, resulting in a significant decrease in the input-output function of L2/3 pyramidal neurons. This is reminiscent of previous findings on the activity-dependent maintenance of intrinsic excitability in L5 pyramidal neurons, where visual deprivation drives a reduction rather than an increase in intrinsic excitability, a mechanism that is also strongly developmentally regulated (Nataraj et al., 2010; Nataraj and Turrigiano, 2011). Notably, the decrease in L2/3 pyramidal neuron intrinsic excitability induced by conjoint reduction of cholinergic inputs and V1 activity would be expected to reduce firing, making this mechanism a likely contributor to the block of upward firing rate homeostasis.

Taken together with previous studies, our data suggest that the cellular mechanisms underlying upward firing rate homeostasis vary as a function of developmental age and likely brain region and cell type. This is consistent with a large literature showing that distinct homeostatic mechanisms are induced at different developmental ages, although in some cases direct comparisons are complicated by the use of different deprivation paradigms (Desai et al., 2002; Keck et al., 2013; Barnes et al., 2015; Keck et al., 2017; Wen and Turrigiano 2021). This complexity suggests that there are likely additional targets of cholinergic neuromodulation within V1 that contribute to the induction of firing rate homeostasis during AW.

HDB ACh axons project to and terminate within V1 (Kim et al., 2016, Figure 1), but also have other targets, including the hippocampus and other cell types within the BF, which also project to V1 (Bloem et al., 2014; Kim et al., 2015; Zant et al., 2016). We therefore wondered whether these inputs directly modulate L2/3 pyramidal neurons via cholinergic receptor expression, or whether the effects of HDB inhibition are mediated indirectly via other targets. To begin to address this, we knocked down the M1 ACh receptor, which is the most abundant ACh receptor in V1, and is expressed at high levels in pyramidal neurons (Levey et al., 1991; Yamasaki et al., 2010; Groleau et al., 2015). We were able to demonstrate that viral-mediated knockdown of M1 within V1 abolished the effects of CCh wash-in on the after-depolarization in L2/3 pyramidal neurons measured *ex vivo*; this effect is known to be mediated by M1 ACh receptors (Gulledge et al., 2009), indicating that knockdown was effective. If this receptor mediates the ability of HDB ACh inputs to maintain intrinsic excitability during V1 activity suppression, then this knockdown should phenocopy inhibition of HDB ACh neurons. In contrast, we found that intrinsic excitability was unchanged by the conjunction of M1 knockdown and activity suppression.

Taken together, our data rule out two possible mechanisms via which ACh could be mediating its gating effects on upward firing rate homeostasis. First, inhibited HDB ACh neurons did not affect how much time animals spent in AW behavioral states, indicating that reduced ACh release from HDB does not gate firing rate homeostasis through brain-wide changes in behavioral state. Second, the ability of HDB ACh neurons to maintain intrinsic excitability is not mediated through M1 AChR knockdown. In these experiments, we targeted our recordings to L2/3 pyramidal neurons that expressed our knockdown construct, but there was robust viral expression in both NeuN^+^ (pyramidal) and NeuN^-^ (non-pyramidal) neurons, indicating that we achieved a wide-spread reduction in M1 within the V1 circuit. Thus, these data suggest that ACh does not work through local M1 receptors in V1 to gate homeostatic plasticity. This leaves open the possibilities that HDB ACh terminals mediate these effects through other receptor subtypes within V1, or that the impact on V1 plasticity is indirect and is mediated through a structure outside of V1, which then targets V1 to gate plasticity. Regardless of the exact targets, our data show that cholinergic signaling during AW plays a critical role in enabling homeostatic increases in V1 excitability during sensory or activity deprivation.

## Acknowledgements

This research was supported by NIH Grant 1F31EY031602-01 (JB) and NIH RO1 EY025613 (GGT). In vivo spike extraction and initial processing were performed using Brandeis University’s High Performance Computing Cluster which is partially funded by DMR-MRSEC 1420382. We thank Alejandro Torrado Pacheco for invaluable mentorship and his work on the *in vivo* recording and analysis pipeline, Daniel Leman for *in vivo* experiment support, Lauren Tereshko for the empty vector control virus, and SciDraw for the rat illustration.

